# Revolutionising Public Health Reference Microbiology using Whole Genome Sequencing: *Salmonella* as an exemplar

**DOI:** 10.1101/033225

**Authors:** Philip Ashton, Satheesh Nair, Tansy Peters, Rediat Tewolde, Martin Day, Michel Doumith, Jonathan Green, Claire Jenkins, Anthony Underwood, Catherine Arnold, Elizabeth de Pinna, Tim Dallman, Kathie Grant

## Abstract

Advances in whole genome sequencing (WGS) platforms and DNA library preparation have led to the development of methods for high throughput sequencing of bacterial genomes at a relatively low cost (Loman et al. 2012; Medini et al. 2008). WGS offers unprecedented resolution for determining degrees of relatedness between strains of bacterial pathogens and has proven a powerful tool for microbial population studies and epidemiological investigations (Harris et al. 2010; Lienau et al. 2011; Holt et al. 2009; Ashton, Peters, et al. 2015). The potential utility of WGS to public health microbiology has been highlighted previously (Köser et al. 2012; Kwong et al. 2013; Reuter et al. 2013; Joensen et al. 2014; Nair et al. 2014; Bakker et al. 2014; D’Auria et al. 2014). Here we report, for the first time, the routine use of WGS as the primary test for identification, surveillance and outbreak investigation by a national reference laboratory. We present data on how this has revolutionised public health microbiology for one of the most common bacterial pathogens in the United Kingdom, the *Salmonellae.*

**DATA SUMMARY:** 1. PHE Salmonella sequencing data is deposited in the Sequence Read Archive in BioProject PRJNA248792.

**IMPACT STATEMENT:** The first human genome cost around $3 billion, and took around 10 years to complete. Advances in DNA sequencing technology (also referred to as whole genome sequencing (WGS)) allow the same feat to be accomplished today for less than $10000 and less than 2 weeks. This remarkable improvement in technology has also led to a step change in microbiology, increasing our understanding of the evolution of major human pathogens such as *Yersinia pestis, Salmonella* Typhi and *Mycobacterium tuberculosis.* While these kinds of academic studies provide unparalleled context for public health action, until now, this approach has not been routinely employed at the frontline. At Public Health England, WGS has been implemented for routine public health identification, characterisation and typing of an important human pathogen, *Salmonella,* replacing methods that have changed little over the last 100 years. Analysis of WGS data has identified outbreaks that were previously undetectable and been used to infer rare antimicrobial resistance patterns. This paper will serve as a notification to the community of the methods PHE are using, and will be of great use to other public health labs considering switching to WGS.

## INTRODUCTION

Governments have a long history of intervening on behalf of the public health. In the 19^th^ Century public health initiatives sprang up to combat the main afflictions of the day, which were primarily microbiological. In the UK, Edwin Chadwick spearheaded a movement that resulted in the passage of the Public Health Act 1848. One of Chadwick’s primary goals was to disperse the ‘miasma’, or polluted air, that was then held cause diseases such as cholera and chlamydia. This was to be achieved by draining the 30 000 cesspools of London into the river Thames. Unfortunately, the Thames was also the primary source of drinking water for the city at that time. Thus, one of the first modern public health interventions contributed to a dramatic increase in cholera rates in the city (Johnson 2006). While John Snow, Louis Pasteur and Robert Koch soon placed microbiological public health on a firmer footing than ‘miasma’, the dangers of basing public health action on inaccurate, or out of date, science remain.

Here, we present the experience of a national reference lab that has undergone a transformation from a traditional serotyping laboratory to a state of the art whole genome sequencing laboratory. As we are the first national reference laboratory to do this for *Salmonella,* we believe that relating our experience and approach will be valuable for the wider community.

### Traditional approach to reference microbiology for *Salmonella* species

Approximately 10,000 *Salmonella* isolates are referred to the Gastrointestinal Bacterial Reference Unit (GBRU), Public Health England (PHE) each year. Prior to April 2015, all presumptive *Salmonella* isolates received were speciated and sub-speciated using real-time PCR (RT-PCR) (Katie L. Hopkins et al. 2011) and phenotypic arrays (Omnilog). Serological classification, as described in the White-Kauffman-Le Minor scheme (Grimont & Weill 2008), utilises the phenotypic variation seen in flagellar, polysaccharide and capsular antigens, was then used to provide further resolution. *Salmonella* resolves into more than 2600 serotypes according to their antigenic formulae and the procedure relies upon the production of numerous antisera raised in rabbits following a precise immunisation programme. The incidence of different *Salmonella* serotypes identified in the UK is not uniform with only 10 serotypes accounting for approximately 70% of the isolates received by GBRU (Figure 1). Serotyping did not always provide the level of strain discrimination required for outbreak investigation, and was complimented with phage-typing (Callow 1959) and reactive (i.e. not routine for all isolates) molecular methods such as Multi Locus Variable number of tandem repeats Analysis (MLVA) (K. L. Hopkins et al. 2011) or Pulsed-Field Gel Electrophoresis (PFGE) (Peters 2009).

**Figure 1:**
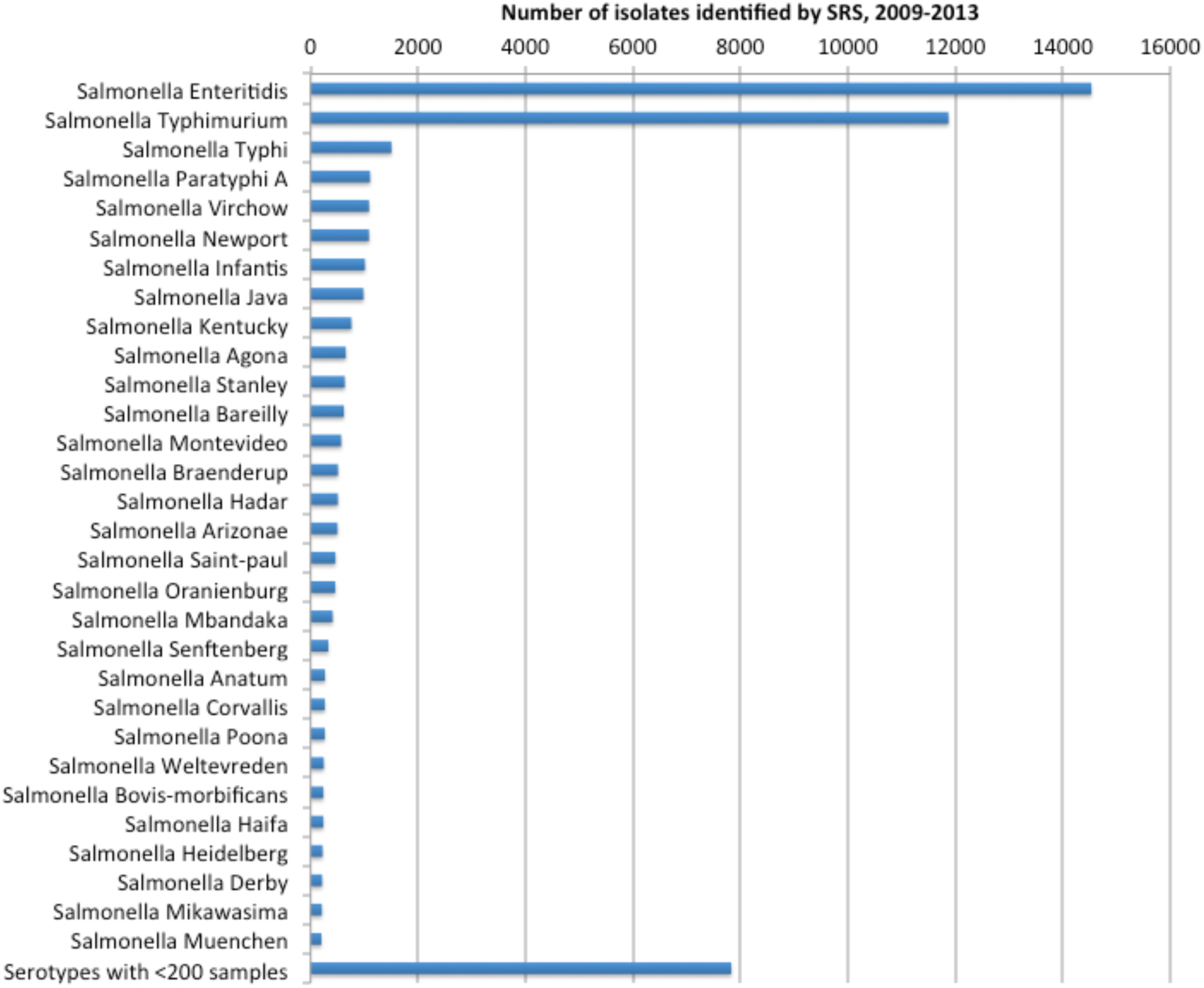
Serotypes with more than 200 isolates received in the 4 year period 2009–2013 by Salmonella Reference Service, Public Health England.

### WGS approach to reference microbiology for *Salmonella* species

#### Genome sequencing

Since April 2015, WGS has been the one procedure performed on all cultures of *Salmonella sp.* referred to GBRU by frontline hospital microbiology laboratories, private laboratories and food, water and environmental laboratories. All other typing methods previously employed have been significantly reduced or withdrawn. On receipt, original cultures are directly inoculated into 750 μl of nutrient broth and incubated over night at 37^o^C. DNA extraction is performed using the Qiasymphony automated DNA extractor (Qiagen) and DNA is quantified using the Glomax (Promega). DNA is submitted to the central Genomic Sequencing and Development Unit at PHE, where Illumina Nextera XT DNA libraries are constructed and sequenced using the Illumina HiSeq 2500 in fast mode. The samples are then deplexed by Casava software and Trimmomatic (Bolger et al. 2014) used to trim any data with a phred score less than Q30 from the beginning and end of the reads. The outline of this process can be seen in Figure 2.

**Figure 2:**
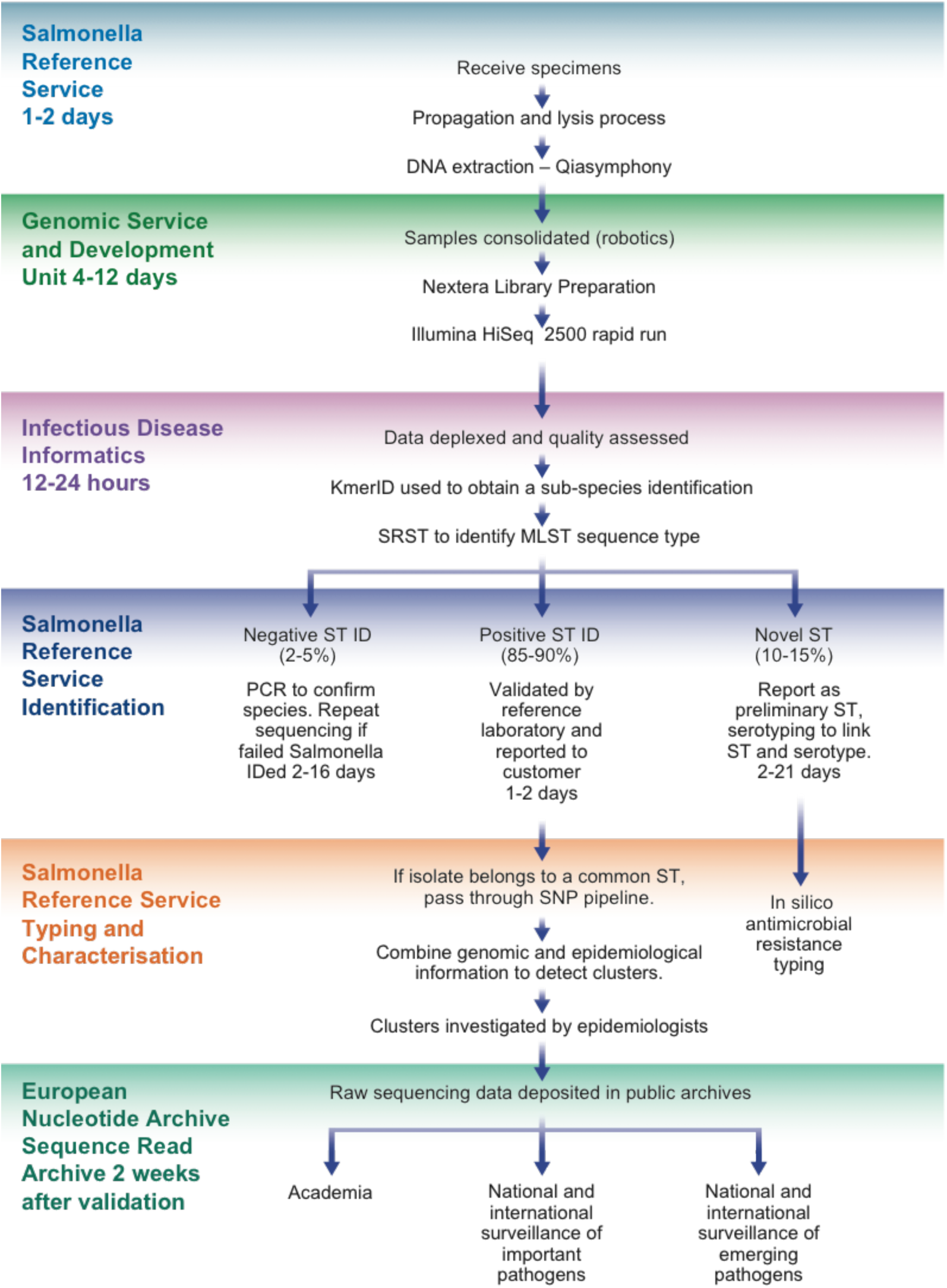
The current workflow for the use of whole genome sequencing by the Public Health England Salmonella Reference Service

#### Bacterial identification and serogroup designation

A sample of k-mers (DNA sequences of length k) in the sequence data are compared against the k-mers of 1769 reference genomes representing 59 pathogenic genera obtained from RefSeq. The closest percentage match is identified, and provides initial confirmation of the species and sub-species of *Salmonella.* This step also identifies samples containing more than one species of bacteria (i.e. mixed cultures) and any bacteria misidentified as *Salmonella* by the sending laboratory.

Once the sample is confirmed as *Salmonella,* the sequenced genome is interrogated with sequences in the Multi Locus Sequence Typing (MLST) database (Achtman et al. 2012) using a modified version of SRST (Inouye et al. 2012). This provides a quality assessed sequence type (ST). Achtman *et al* have shown that *Salmonella* serotypes generally belong to clonal complexes of related Sequence Types (ST) known as e-burst groups (EBGs) and have described the correlation between EBGs and serotype (Achtman et al. 2012). At PHE a database containing matched MLST data and phenotypic serotype designation for more than 12000 isolates of *Salmonella* allows us to infer a serogroup from the sequence type with 96% accuracy (Ashton, Nair, et al. 2015). The primary error types are (i) two different serotypes having the same ST, e.g. ST 909 is both Salmonella Richmond and Salmonella Bareilly (ii) processing errors and (iii) inaccuracy in serotype designation of public data (Ashton, Nair, et al. 2015). The inferred serotype is then reported back to the sending laboratory, along with the ST of the isolate, to maintain backwards compatibility with historical data. This provides customers with a consistent service and facilitates data exchange with Public Health colleagues both locally and internationally, as well as others in the veterinary, food, water and environmental microbiology disciplines.

#### Outbreak detection and investigation

Over 70% of isolates received by GBRU belong to the most common 14 serotypes and, as with serotype, ST alone is not discriminatory enough for outbreak detection and public health investigation. Whole genome single nucleotide polymorphism (SNP) typing is performed on samples that belong to the most common e-burst groups. This involves mapping the sequence reads to an appropriate reference genome (within the same e-burst group) using BWA mem (Li & Durbin 2009), before identifying SNPs with GATK (DePristo et al. 2011). High quality SNPs are then stored in a database where they can be queried to generate sequence alignments for input into phylogenetic algorithms.

Within PHE, outbreaks of *Salmonella* were traditionally detected using an exceedance algorithm (Noufaily et al. 2013) based on serotyping and phage typing data. The lack of discrimination associated with these typing techniques results in a high false positive rate of exceedance notifications (Noufaily et al. 2013). In contrast, whole genome SNP typing offers unprecedented resolution in linking cases providing added certainty to outbreak definitions, transmission networks and other aspects of the underlying epidemiology.

A hierarchical ‘SNP address’ approach is employed that groups isolates together into clusters of increasing levels of similarity. The pairwise SNP distance is calculated for each pair of isolates in the analysis set. This distance matrix is then subjected to single linkage clustering at 250, 100, 50, 25, 10, 5 and 0 SNPs. The end result is a SNP address that identifies clusters of isolates at each level of the hierarchy (Figure 3). The SNP address approach for identifying epidemiologically significant clusters (i.e. outbreaks) correlates well with existing workflows and is phylogenetically informative, as all isolates are placed into haplotypes derived from a phylogeny of the clonal complex in question.

**Figure 3:**
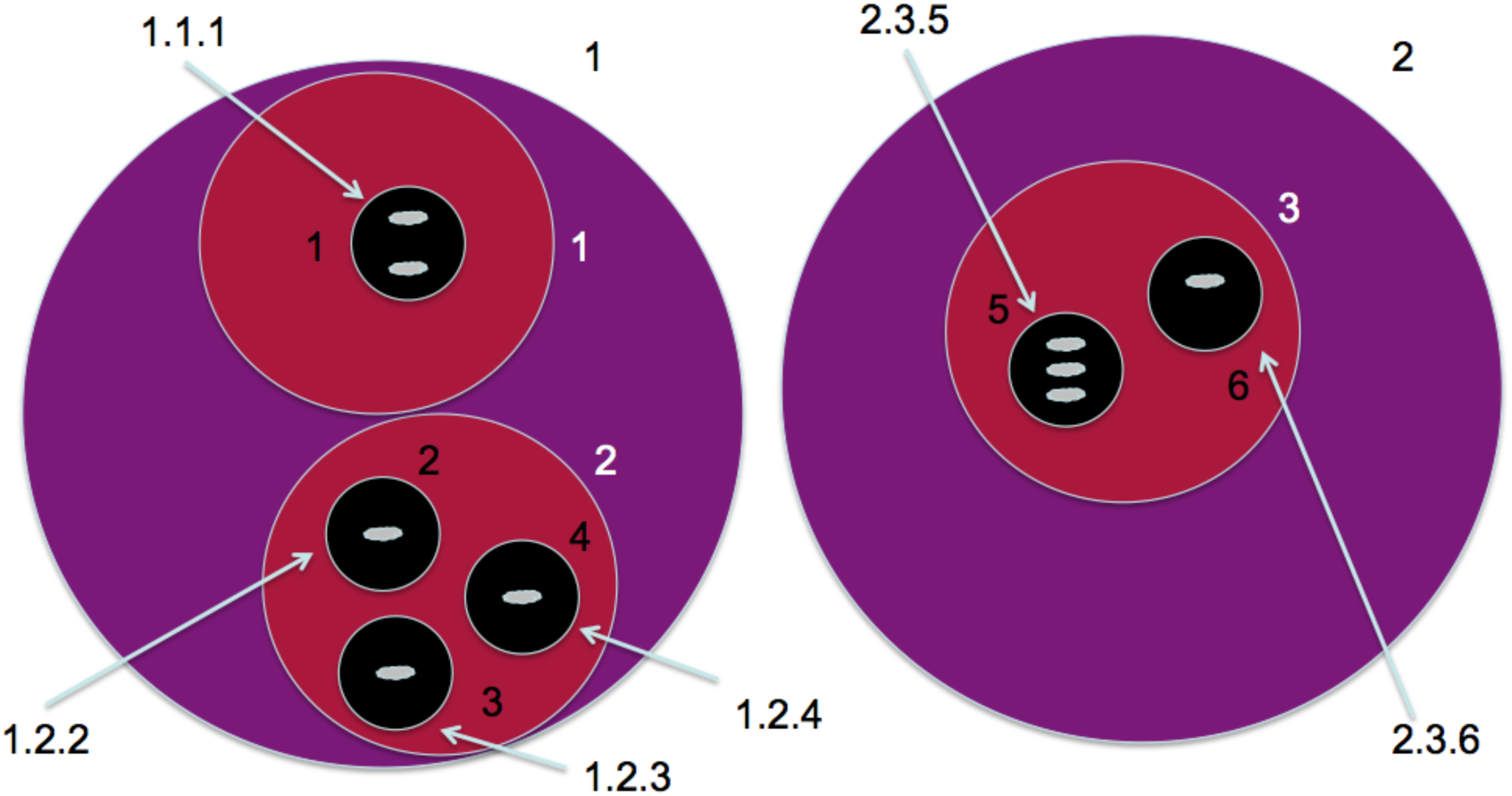
Schematic of how the SNP address works

Median turn around time from the receipt of a sample at the reference laboratory to a reported, WGS based, result over the last 4 weeks before submission of this manuscript (weeks 43–46, 2015) months was 10 working days (David Powell, PHE, personal communication). Once an outbreak has been identified it is important to confirm additional cases quickly in order to expedite epidemiological investigations. To this end we are exploring the use of WGS data to design outbreak/incident-specific RT-PCR or High Resolution Melt assays to screen additional referred cultures as being part of specific outbreaks based on SNPs that are unique to this cluster. Then, in the case of an incident, all samples could be rapidly screened upon receipt to ascertain if they are part of the outbreak. Epidemiological investigation could then proceed in a more timely fashion. These isolates could be sequenced as a matter of urgency using a MinION device (Oxford Nanopore Technologies), for example, which could confirm the isolate as being part of a known outbreak, or not (Quick et al. 2015) (PMID 26025440).

#### Further microbiological characterisation

One of the most attractive aspects of WGS is the fact that it lends itself to a ‘single process, multiple tests’ approach. For example, once you have generated the sequence in order to do whole genome

SNP typing, the data is available in perpetuity for other tests, e.g. characterisation of virulence and other molecular markers e.g. of antimicrobial resistance. Known antimicrobial resistance determinants can be readily identified from WGS data. We carried out a pilot study to compare genotypic vs phenotypic resistance typing in 642 Salmonellae of which 57.5% were susceptible and 24.7% MDR. Results showed a greater than 99% success rate (unpublished data) indicating the potential of adding this into the routine *Salmonella* work flow. Phenotypic screening could be used on a small proportion of isolates in order to detect novel or emerging resistance mechanisms. The fact that the data is available in perpetuity allows re-testing when e.g. novel resistance mechanisms emerge.

#### Data sharing

The genomic data of all *Salmonella* sequenced at PHE is publically released into the NCBI BioProject PRJNA248792 within two weeks of the sample report date. The prompt release of data is to facilitate international tracking and surveillance of food and waterborne, gastrointestinal pathogens in far-reaching distribution networks (Byrne et al. 2015). Globally, national surveillance organizations, such as the Food and Drug Administration and the United States Centre for Disease Control, are also uploading *Salmonella* genomes from their surveillance activities into public sequence archives. Algorithms that enable the timely and sensitive comparison of these datasets are a high priority, as this will be needed to monitor international patterns in gastrointestinal infection. While meta-data is limited for privacy reasons, publically releasing the data provides a great resource for academic researchers interested in gastrointestinal pathogens.

#### Infrastructure requirements and challenges

WGS is obviously a very exciting technology, however it is one that requires substantial investment in infrastructure. Public Health England have invested millions in sequencing and molecular biology hardware, high performance computing and staff costs (wet lab and bioinformaticians) to deliver this service. There are also many challenges involved in changing skillsets and mindsets. Practical implementation and integration across different departments (microbiology, sequencing, bioinformatics and epidemiology) is also a challenge, but necessary to ensure the success of a project such as this.

### Future perspectives

It is an exciting time to work in microbiology and genomics. Even before the repercussions of one revolution (short read, whole genome sequencing) have had their full impact on reference and clinical microbiology, another revolution (long read, portable sequencing) is on the horizon. Exactly how this second revolution will affect clinical and reference microbiology is currently unclear but two obvious applications present themselves. Firstly, the ability to fully assemble large numbers of genomes could provide a step-change in resolution for determining whether two genomes are related. Currently, it is the core genome that is the focus of the majority of genomic epidemiology. Being able to analyse the entire genome provides opportunities for using the accessory genome to determine how related two isolates are. However, this approach needs to be thoroughly investigated and assessed for sensitivity and specificity as compared with a core genome approach. We need to be mindful that the dynamic accessory genome could be misleading as to the relatedness of isolates (Lauren Cowley, PHE, personal communication). The other application of this new wave of sequencers comes from their portability and low profile infrastructure requirements. Many smaller hospitals and front line laboratories could not justify setting up the infrastructure required to run a large machine as they are unlikely to perform enough sequencing to make it economically viable. However, it would be feasible to employ smaller devices like the MinION and use them to sequence suspect outbreak strains. The data could be streamed to a repository maintained by the reference laboratory where the SNPs identified in the sample could be used to place the sample onto a tree and call it as outbreak or non-outbreak (Figure 4). This approach has already been used in an outbreak situation (Quick et al. 2015). There is a big question for many organisations as to whether to invest heavily now in proven, ‘work-horse’ machines, or to hold on for exciting new technologies to mature, with the potential increases in throughput and decrease in cost promised by these ‘third generation’, or ‘third revolution’ (Loman & Pallen 2015), technologies.

**Figure 4:**
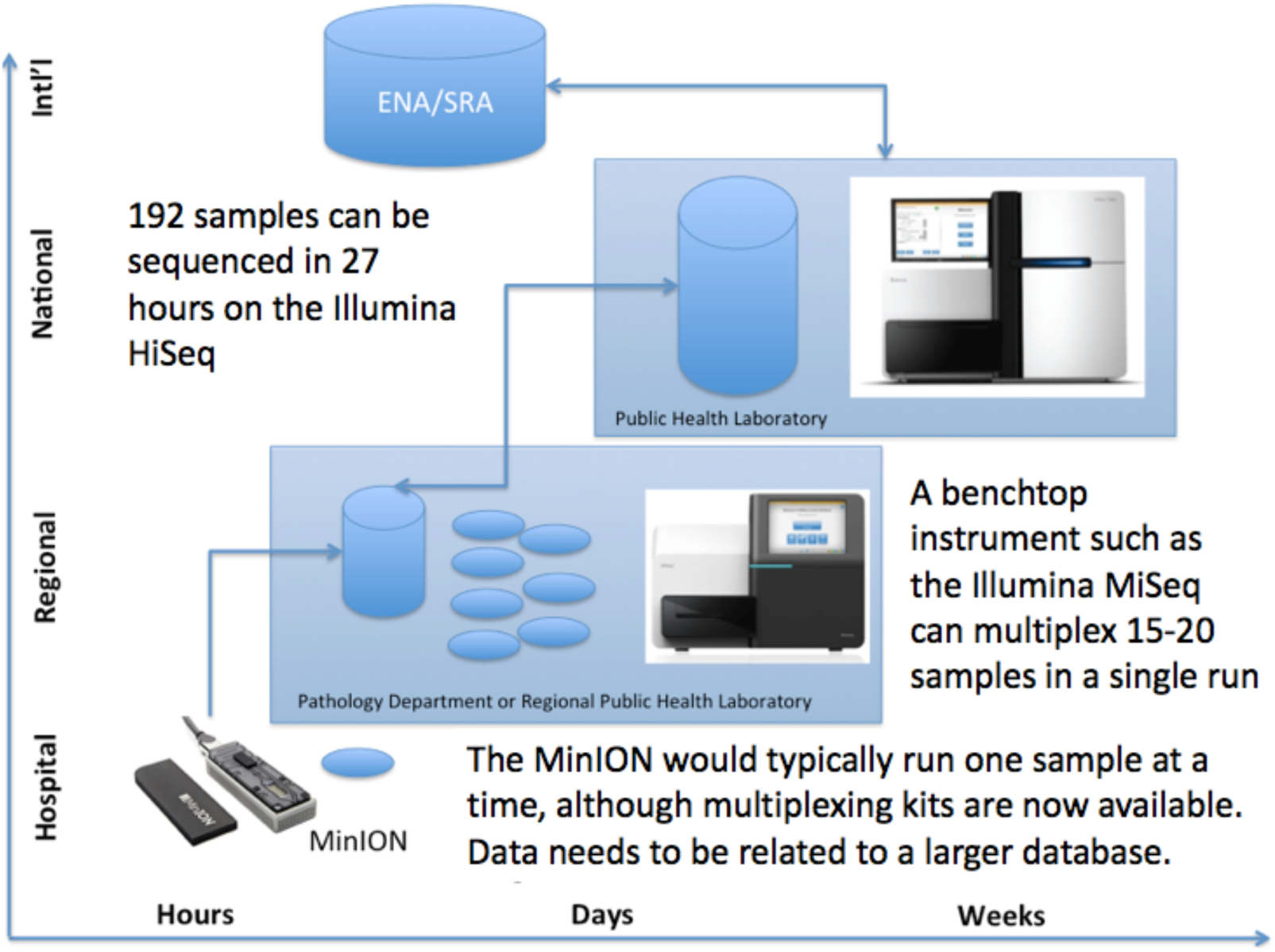
Example of future public health workflow

## CONCLUSION

WGS for routine public health surveillance offers a host of scientific, technical and economic advantages compared with conventional microbiological methods. A single, highly automatable process provides speciation, identification, typing and characterization to a level at least equivalent, and often superior, to the previous ‘gold-standard’ techniques. In addition, the data is uniquely portable, allowing rapid comparison with other public health institutions and re-use by the wider scientific community.

## ACKNOWLEDGEMENTS

We would like to acknowledge the contribution of everyone working in the Salmonella Reference Service, the Genome Sequencing and Development Unit and Infectious Disease Informatics at Public Health England, for their invaluable contributions to this work. We would also like to thank Nick Loman for significant contribution toward Figure 4.

## REFERENCES

Achtman, M. et al., 2012. Multilocus Sequence Typing as a Replacement for Serotyping in Salmonella enterica. PLoS pathogens, 8(6), p.e1002776. Available at: http://www.pubmedcentral.nih.gov/articlerender.fcgi?artid=3380943&tool=pmcentrez&rendertype=abstract [Accessed July 20, 2012].

Ashton, P.M., Nair, S., et al., 2015. Identification and typing of Salmonella for public health. Peer J Pre Prints, 3(April), p.e1778.

Ashton, P.M., Peters, T., et al., 2015. Whole Genome Sequencing for the Retrospective Investigation of an Outbreak of Salmonella Typhimurium DT 8. PLoS Currents. Available at: http://currents.plos.org/outbreaks/?p=46522.

Bakker, H.C. Den et al., 2014. Rapid Whole-Genome Sequencing for Surveillance of Salmonella enterica Serovar Enteritidis. Emerging infectious diseases, 20(8), pp.1306–14. Available at: http://www.pubmedcentral.nih.gov/articlerender.fcgi?artid=4111163&tool=pmcentrez&rendertype=abstract.

Bolger, A.M., Lohse, M. & Usadel, B., 2014. Trimmomatic: a flexible trimmer for Illumina sequence data. Bioinformatics (Oxford, England), pp.1–7. Available at: http://www.ncbi.nlm.nih.gov/pubmed/24695404.

Byrne, L. et al., 2015. A multi-country outbreak of Salmonella Newport gastroenteritis in Europe associated with watermelon from Brazil, confirmed by whole genome sequencing: October 2011 to January 2012. Euro surveillance: bulletin Européen sur les maladies transmissibles = European communicable disease bulletin, 19(31), pp.6–13.

Callow, B.R., 1959. A new phage-typing scheme for Salmonella Typhimurium. The Journal of hygiene, 57, pp.346–359.

D’Auria, G., Schneider, M.V. & Moya, A., 2014. Live genomics for pathogen monitoring in public health. Pathogens (Basel, Switzerland), 3(1), pp.93–108. Available at: http://www.pubmedcentral.nih.gov/articlerender.fcgi?artid=4235738&tool=pmcentrez&rendertype=abstract.

DePristo, M.A. et al., 2011. A framework for variation discovery and genotyping using next-generation DNA sequencing data. Nature …, 43(5), pp.491–498. Available at: http://www.nature.com/ng/journal/vaop/ncurrent/full/ng.806.html [Accessed January 6, 2014].

Grimont, P. & Weill, F.-X., 2008. Antigenic formulae of the Salmonella servovars, WHO Collaborating Centre for Reference and Research on Salmonella. Available at: http://www.pasteur.fr/ip/portal/action/WebdriveActionEvent/oid/01s-000036-89\npapers2://publication/uuid/CA3447A0-61BF-4D62-9181-C9BA78A.

Harris, S.R. et al., 2010. Evolution of MRSA During Hospital Transmission and Intercontinental Spread., 327(5964), pp.1–11.

Holt, K.E. et al., 2009. High-throughput sequencing provides insights into genome variation and evolution in Salmonella Typhi. Nature Genetics, 40(8), pp.987–993.

Hopkins, K.L. et al., 2011. A novel real-time polymerase chain reaction for identification of Salmonella enterica subspecies enterica. Diagnostic Microbiology and Infectious Disease, 70(2), pp.278–280. Available at: http://dx.doi.org/10.1016/j.diagmicrobio.2011.01.015.

Hopkins, K.L. et al., 2011. Standardisation of multilocus variable-number tandemrepeat analysis (MLVA) for subtyping of Salmonella enterica serovar Enteritidis. Eurosurveillance, 16(32), pp.1–11.

Inouye, M. et al., 2012. Short read sequence typing (SRST): multi-locus sequence types from short reads. BMC genomics, 13(1), p.338. Available at: http://www.pubmedcentral.nih.gov/articlerender.fcgi?artid=3460743&tool=pmcentrez&rendertype=abstract [Accessed March 9, 2013].

Joensen, K.G. et al., 2014. Real-time whole-genome sequencing for routine typing, surveillance, and outbreak detection of verotoxigenic Escherichia coli. Journal of clinical microbiology, 52(5), pp.1501–10. Available at: http://www.pubmedcentral.nih.gov/articlerender.fcgi?artid=3993690&tool=pmcentrez&rendertype=abstract [Accessed May 27, 2014].

Johnson, S., 2006. The Ghost Map, Tantor Media, Inc.

Köser, C.U. et al., 2012. Routine Use of Microbial Whole Genome Sequencing in Diagnostic and Public Health Microbiology G. F. Rall, ed. PLoS Pathogens, 8(8), p.e1002824. Available at: http://dx.plos.org/10.1371/journal.ppat.1002824 [Accessed August 3, 2012].

Kwong, J.C. et al., 2013. Whole genome sequencing in diagnostic and public health microbiology. Pathology, 47(May), p.2013.

Li, H. & Durbin, R., 2009. Fast and accurate short read alignment with Burrows-Wheeler transform. Bioinformatics (Oxford, England), 25(14), pp.1754–60. Available at: http://www.pubmedcentral.nih.gov/articlerender.fcgi?artid=2705234&tool=pmcentrez&rendertype=abstract [Accessed August 6, 2013].

Lienau, E. et al., 2011. Identification of a salmonellosis outbreak by means of molecular sequencing. … England Journal of…, 364, pp.981–982. Available at: http://www.nejm.org/doi/full/10.1056/NEJMc1100443 [Accessed July 5, 2013].

Loman, N.J. et al., 2012. Performance comparison of benchtop high-throughput sequencing platforms. Nature Biotechnology, 2012(March). Available at: http://www.nature.com/doifinder/10.1038/nbt.2198 [Accessed April 22, 2012].

Loman, N.J. & Pallen, M.J., 2015. Twenty years of bacterial genome sequencing. Nature Reviews Microbiology, (November), pp.1–9. Available at: http://www.nature.com/doifinder/10.1038/nrmicro3565.

Medini, D. et al., 2008. Microbiology in the post-genomic era. Nature reviews. Microbiology, 6(6), pp.419–430.

Nair, S. et al., 2014. Salmonella enterica subspecies II infections in England and Wales - The use of multilocus sequence typing to assist serovar identification. Journal of Medical Microbiology, 63(PART 6), pp.831–834.

Noufaily, A. et al., 2013. An improved algorithm for outbreak detection in multiple surveillance systems. Statistics in medicine, 32(7), pp.1206–22. Available at: http://www.ncbi.nlm.nih.gov/pubmed/22941770 [Accessed June 6, 2014].

Peters, T.M., 2009. Pulsed-Field Gel Electrophoresis for Molecular Epidemiology of Food Pathogens. In pp. 59–70. Available at: http://link.springer.com/10.1007/978-1-60327-999-4_6.

Quick, J. et al., 2015. Rapid draft sequencing and real-time nanopore sequencing in a hospital outbreak of Salmonella. Genome Biology, 16(1), pp.1–14. Available at: http://genomebiology.com/2015/16/1/114.

Reuter, S. et al., 2013. Rapid bacterial whole-genome sequencing to enhance diagnostic and public health microbiology. JAMA internal medicine, 173(15), pp.1397–404. Available at: http://www.pubmedcentral.nih.gov/articlerender.fcgi?artid=4001082&tool=pmcentrez&rendertype=abstract.

